# Sortase-mediated site-specific modification of interleukin-2 for the generation of a tumor targeting acetazolamide-cytokine conjugate

**DOI:** 10.1101/2020.07.20.211870

**Authors:** Baptiste Gouyou, J Millul, A Villa, S Cazzamalli, D Neri, M Matasci

## Abstract

1.

Small ligands specific to tumor-associated antigens can be used as alternatives to antibodies for the delivery of small payloads such as radionuclides, cytotoxic drugs and fluorophores. Their use as delivery moiety of bioactive proteins like cytokines remains largely unexplored. Here, we describe the preparation and *in vivo* characterization of the first small molecule-cytokine conjugate targeting carbonic anhydrase IX (CAIX), a marker of renal cell carcinoma and hypoxia. Site-specific conjugation between interleukin-2 and acetazolamide was obtained by Sortase A-mediated transpeptidation. Binding of the conjugate to the cognate CAIX antigen was confirmed by surface plasmon resonance. The *in vivo* targeting of structures expressing carbonic anhydrase IX was assessed by biodistribution experiments in tumor bearing mice. Optimization of manufacturability and tumor targeting performance of acetazolamide-cytokine products will be required in order to enable industrial applications.

**Graphical abstract:** 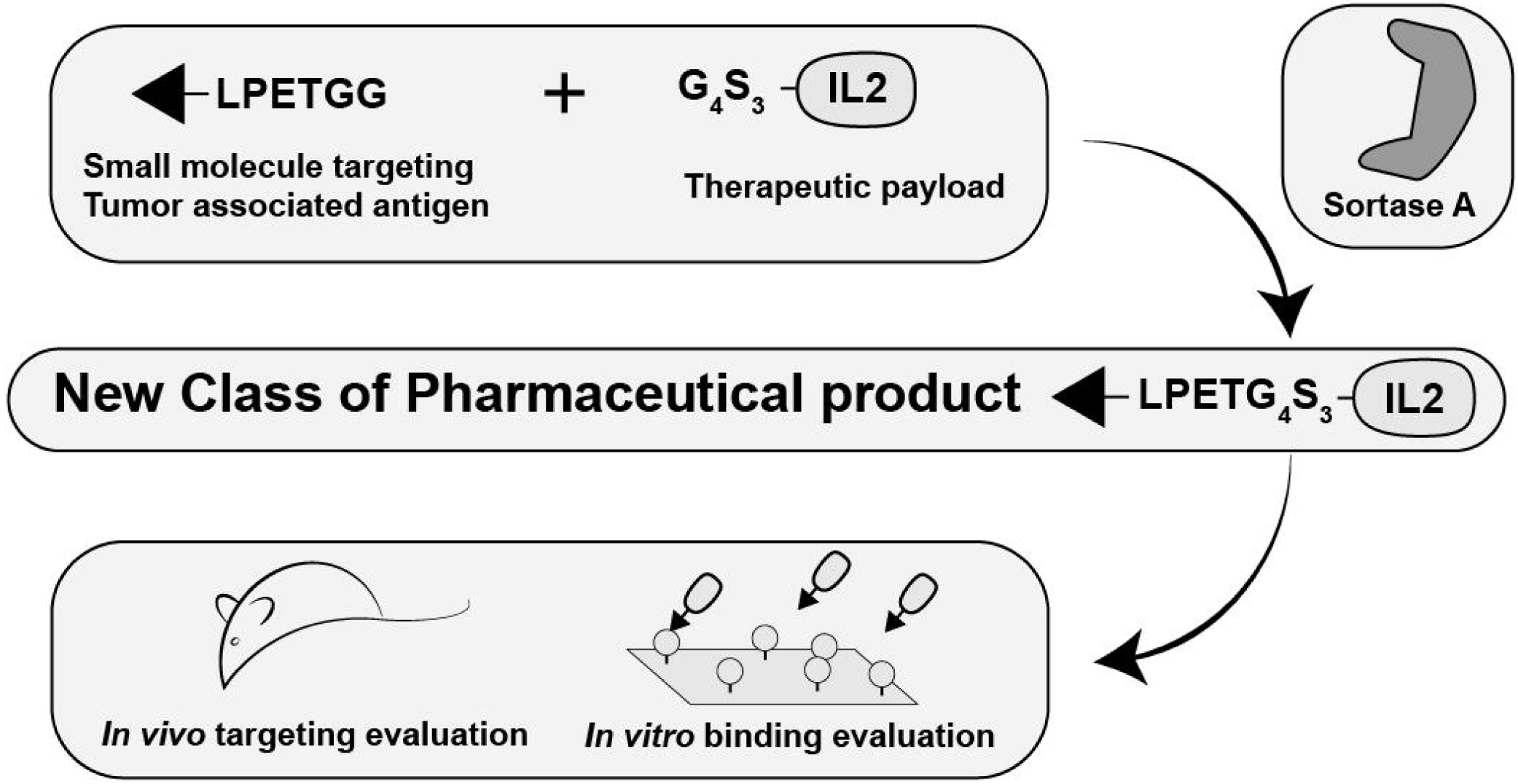

## 2. Introduction

The active delivery of anticancer cytokines to tumor tissue has been extensively explored with the aim to enhance their therapeutic window and circumvent adverse events associated with their systemic administration. To this respect tumor specific antibodies have been used as selective vehicles for the targeted delivery of pro-inflammatory cytokines including interleukin-2 (IL2), intereleukin-12 (IL12) and tumor necrosis factor (TNF)^1–6^. In several cases antibody-cytokine fusion proteins (i.e. Immunocytokines), have demonstrated selective tumor uptake and anti-tumor activity in murine models of cancer and several products have been advanced into clinical trial ^2,3,6–8^. However, the use of antibodies for pharmacodelivery of bioactive payloads to solid tumors, may be limited by their relatively large size, which can lead to slow extravasation and poor tumor tissue penetration combined with extended systemic halflife^9^. These limitations can be partly counteracted by using small antibody fragments (like scFv or diabodies)^10^.

Small organic ligands specific for a list of tumor-associated antigens (e.g., Folate Receptor, Prostate-Specific Membrane Antigen, Somatostatin Receptors and Carbonic Anhydrase IX (CAIX))^11–14^ have been considered as an alternative to antibodies for the targeted delivery of drugs and of therapeutic radionuclides^15^. Small molecules can diffuse very rapidly and homogeneously into solid tumors, potentially reaching high tumor-to-organ ratios by combining extended residence time at the tumor site with rapid body clearance^12,14^.

CAIX is a cell-surface non-internalizing marker of tumor hypoxia that is highly expressed in about 90% of clear cell renal cell carcinomas (RCCs) and in other aggressive cancers^16^. In contrast, CAIX is virtually absent in most normal adult tissues, exception made for some structures of the gastro-intestinal tract^17^.

Our group has demonstrated that acetazolamide (AAZ), a small molecule binder of CAIX, can be effectively used to deliver radionuclides and cytotoxic drugs to tumors for diagnostic and therapeutic applications in tumor bearing mice^18–20^. We have recently reported good SPECT-CT imaging results in patients with renal cell carcinoma using a derivative of acetazolamide labeled with the radioactive payload 99mTc^21^.

The IL2 cytokine is a potent inducer of cytotoxic T cells and NK cells, and was one of the first immunotherapeutic agents approved by FDA for the treatment of metastatic melanoma and renal cell carcinoma (RCC)^22^. However, its use in RCC at high dose produces durable complete response in a small portion of patients but with severe systemic toxicity^23,24^. The antibody-based delivery to tumors has shown the potential to improve the therapeutic index of IL2 in immunocompetent mouse models of cancer^2,3,5,6^.

Traditional strategies for covalent bioconjugation allow limited control over the site and frequency of the modification, resulting in heterogeneous products with potential loss of biological activity of the modified protein^25^. Sortase A (SrtA) is a sequence-specific transpeptidase that catalyze the ligation between LPXTG-containing polypeptides (Sortag) and oligoglycine terminated moieties^26^. Due to its high specificity for the Sortag and broad substrate tolerance, SrtA-mediated transpeptidation has emerged as a powerful method for the site-specific modification of proteins. Recently SrtA has been used for the site specific conjugation of antibodies to generate products such as imaging probes^26–28^, bispecific antibodies^29^ and antibody drug conjugates^30^.

Here we describe the development and the *in vitro* and *in vivo* characterization of the first small-molecule cytokine conjugate (SMCC) targeting CAIX, termed AAZ–IL2, that was generated by SrtA mediated site specific transpeptidation.

## 3. Results and discussion

The aim of this work was to develop a small molecule-cytokine conjugate (SMCC) targeting CAIX and perform a proof-of-concept study to test its tumor targeting performance *in vivo*. IL2, the model bioactive payload of choice, was coupled to AAZ, a known nanomolar binder of CAIX. The product was generated by SrtA mediated conjugation^26–30^, which allowed to obtain a site-specific functionalization of the IL2 cytokine with the AAZ targeting ligand (**Figure 1A**)

**Figure 1:**
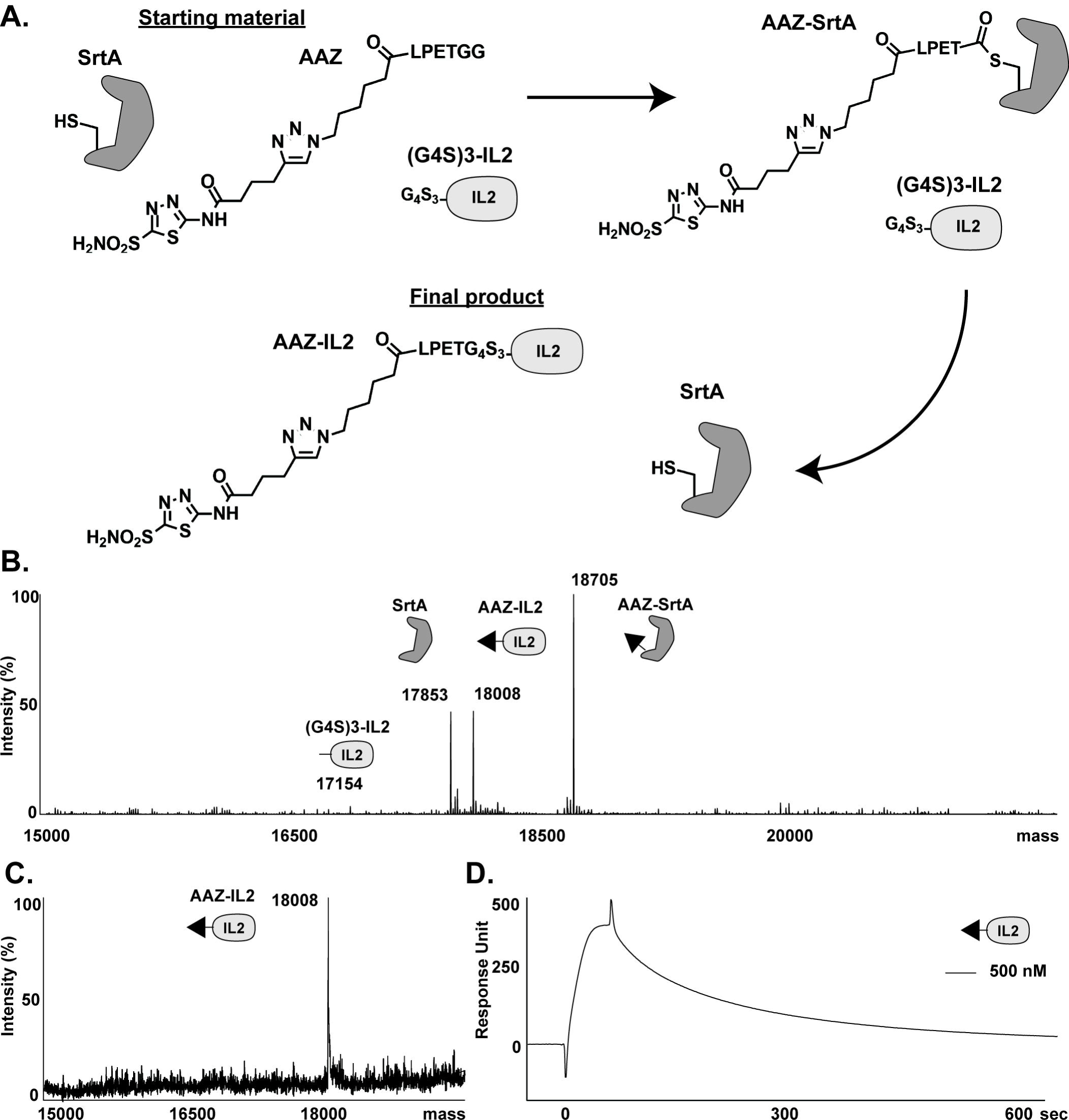
AAZ-IL2 can be conjugated via Sortase A and the pure material is binding to CAIX on *in vitro*. (A) Schematic representation of the Sortase A mediated transpeptidation reaction between AAZ-LPETGG and (G_4_S)_3_-IL2. (B) ESI-MS profile of the crude after 3h of enzymatic reaction. (C) ESI-MS result of AAZ-IL2 after affinity chromatography purification. (D) SPR sensograms of AAZ-IL2 at 500nM on CAIX-coated sensor chip.

The sortagged AAZ-LPETGG substrate was obtained by solid phase peptide synthesis and “on resin” click chemistry of the AAZ moiety, as previously described^14^ followed by standard solid phase peptide synthesis (SPPS) procedures to attach the LPETGG Sortag (**FigureS1A** and **S1B**). Prior to the enzymatic coupling, the identity and purity of the produced AAZ-LPETGG material was analyzed by LC-MS, confirming the presence of a single peak (MW 986 Da) (Figure S1D). The histidine-tagged (G4S)3-IL2 cytokine and SrtA enzyme were recombinantly expressed in mammalian cells and bacteria, respectively, and purified via metal affinity chromatography (Figure S2A S2B). The mutations T23S and C125S were introduced in the (G4S)3-IL2 protein in order to avoid glycosylation and the formation of covalent multimers as confirmed by SDS-PAGE and size exclusion chromatography (Figure S2B).

Complete conjugation of (G4S)3-IL2 to AAZ-LPETGG, could be obtained upon incubation for 3h at 4°C with SrtA, using a 20-fold molar excess of the AAZ-LPETGG substrate. Figure 1B shows the LC-MS analysis of the crude reaction mixture, confirming the virtually complete conversion of the (G4S)3-IL2 substrate (MW 17154 Da) into the final AAZ-IL2 product (MW 18008 Da). Interestingly under these experimental conditions the majority of the SrtA enzyme was still bound to AAZ-LPETGG as covalent thioacyl intermediate (MW 18705 Da) suggesting the possibility for a further optimization of the transpeptidation reaction. The crude reaction product was then subjected to anti-IL2 affinity chromatography to separate the AAZ-IL2 product from unreacted thioacyl intermediate and SrtA. Homogeneity of the purified AAZ-IL2 product was confirmed by LC-MS analysis (**Figure 1C**) and gel filtration chromatography (**Figure S2C**). Furthermore, the capability of AAZ-IL2 to bind CAIX was tested *in vitro* by SPR analysis showing a relatively fast k_on_ and k_off_ (**Figure 1D**), whereas the non-targeted control (G4S)3-IL2 displayed no binding capacity on the CAIX coated chip (data not shown).

SrtA mediated transpeptidation resulted in the production of highly homogeneous site specifically conjugated AAZ-IL2 which retained CAIX *in vitro* binding. However, main limitations in this technology may hamper its application for commercial scale manufacturing, including the need for GMP grade SrtA and reaction substrates, the relatively high amount of SrtA required to achieve complete conversion, and the final purification step necessary to separate the conjugated product from the SrtA enzyme. The use of engineered SrtA variants^31^ with increased catalytic activity, combined with the immobilization of the SrtA to solid supports may contribute to overcome some of these limitations^28^.

Pure preparation of AAZ-IL2 was used to study its quantitative biodistribution in nude mice bearing subcutaneously-grafted human SKRC-52 renal cell carcinomas. The conjugate was radiolabeled with I-125 following previously described methodologies^32^ and injected intravenously at a fixed dose of 0.5 mg/Kg. Mice were sacrificed at different time points and radioactivity in tumors and in different healthy organs was counted. Surprisingly AAZ-IL2 failed to preferentially accumulate at the tumor site at any tested time-point (**Figure 2A**). Furthermore, at the earliest timepoints (up to 6h) a nonnegligible accumulation in the kidney, lung, spleen and stomach was observed, whereas at 24h post-injection all organs were virtually free from AAZ-IL2 (with the exception of some weak uptake into lung and spleen). Uptake into kidney, lung, spleen and stomach seems to be an intrinsic property of the AAZ-IL2 product since a similar behavior was not observed for the untargeted control (**Figure 2B)**. Furthermore, a comparable accumulation in the same normal organs was reported for the biodistribution of 99mTc labelled AAZ derivatives^19^. In these studies, showing efficient AAZ tumor uptake at molecular doses comparable to the ones used in our experiments, targeting selectivity could be improved by increasing the injected dose of the CAIX ligands or by preinjecting unlabeled ligand preparations. It remains to be tested whether a similar approach might be useful to improve the tumor targeting ability of AAZ-IL2.

**Figure 2:**
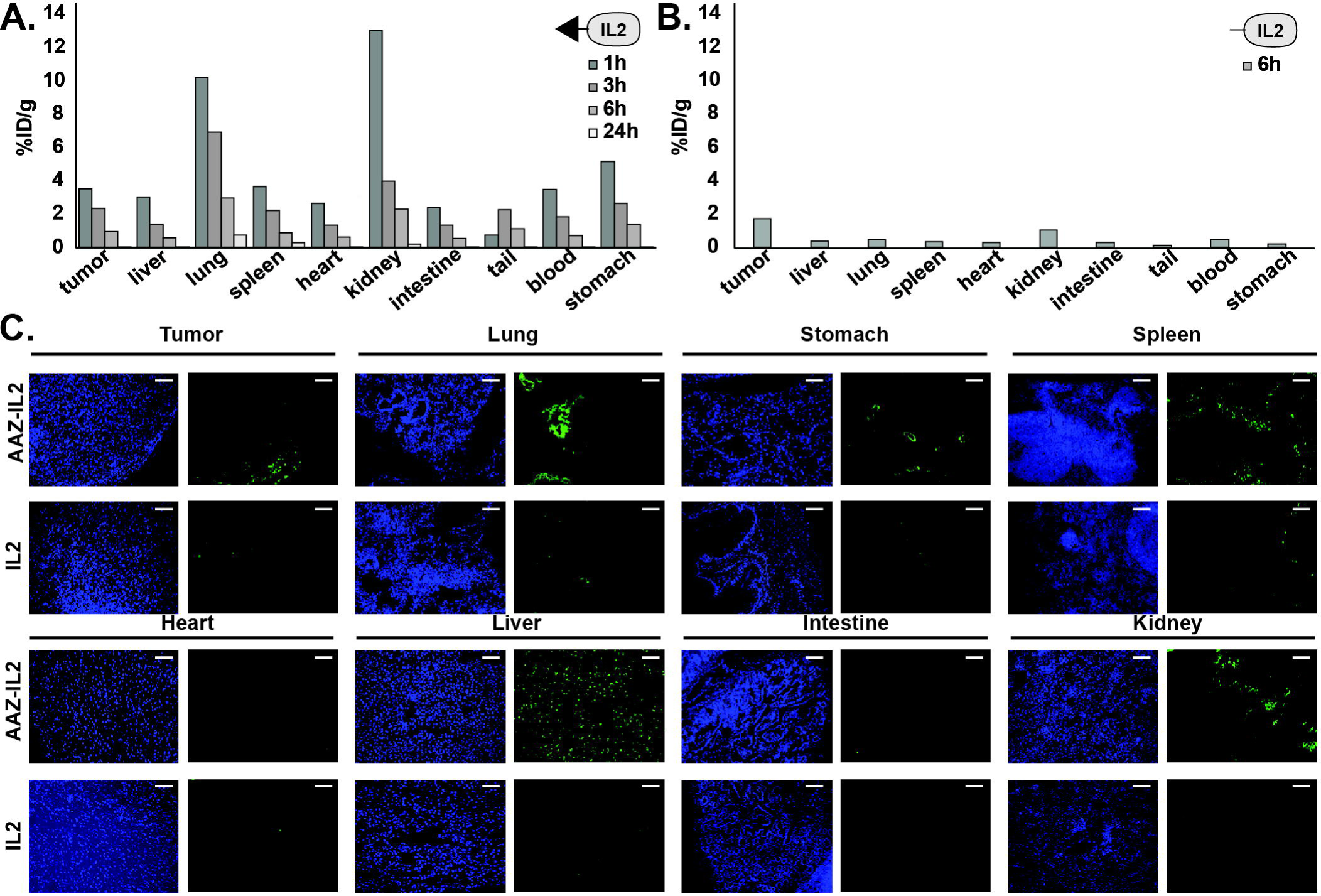
When radiolabeled and fluorescently labeled AAZ-IL2 biodistribution is suboptimal. Quantitative biodistribution analysis of AAZ-IL2 (A) and (G_4_S)_3_-IL2 (B) based fusion proteins. (C) *ex vivo* immuno-fluorescence biodistribution of AAZ-IL2 and (G_4_S)_3_-IL2. The white scale bar corresponds to 50*µ*m, 10x magnification.

A substantial lack of preferential tumor uptake was confirmed in ex-vivo immunofluorescence studies. In this case, tumor bearing mice were sacrificed 2h after the injection of 150 µg of FITC labelled AAZ-IL2 or (G4S)3-IL2 and tissue sections were analyzed for the presence of the injected compounds by immunofluorescence. Only a weak presence of AAZ-IL2 could be detected into tumor sections, whereas a relatively stronger staining was again observed in lung, spleen and kidney (**Figure 2C**). In agreement with the biodistribution results no significant staining was observed for the (G4S)3-IL2 negative control in any of the tissues analyzed.

Our biodistribution studies suggest that AAZ-IL2 reaches the tumor mass already at 1h post injection (earlier time point tested), however no significant specific accumulation could be observed over time. In order to bind the CAIX antigen present on the surface of SKRC-52 cancer cells, AAZ-IL2 needs to efficiently diffuse within the tumor mass. It may be hypothesized that limitations in extravasation and tumor penetration due to the relatively large size (18 KDa) of AAZ-IL2, may impair its targeting properties. Poor tumor tissue penetration and heterogeneous distribution was indeed observed for AAZ-IL2 by *ex vivo* immunofluorescence studies at 2h post injection.

*In vitro* AAZ-IL2 showed binding to CAIX, with a quite low affinity characterized by a fast k_off_. Enhancing tumor uptake and improving tumor to healthy organs selectivity, may be achieved by increasing the affinity to CAIX antigen expressed on the surface of cancer cells.

In the same SKRC-52 xenograft model used in our studies, Krall et al reported that a bivalent-AAZ-dye conjugate with improved CAIX binding affinity, had longer residence on the tumor than the corresponding monovalent version^18^. Similarly, an affinity matured version of acetazolamide recovered from DNA encoded library^33^, showed substantially improved tumor targeting performances when compared to acetazolamide^14,33^. Whether the use of a higher affinity CAIX ligand variant would improve also the targeted delivery of IL2 remains to be elucidated.

## 4. Summary and outlook

To the best of our knowledge, this is the first report of the site-specific conjugation of a small-molecule ligand to a cytokine for tumor targeting purposes. Using SrtA based transpeptidation, we efficiently conjugate the acetazolamide targeting moiety to IL2. This novel small molecule-cytokine conjugate retained binding capacity to CAIX *in vitro*, but lacked tumor targeting specificity *in vivo*, when tested in the SKRC-52 xenograft model of renal cell carcinoma. Whereas further optimization of the conjugate product is required to obtain efficient tumor targeting, including the use of AAZ ligand variant with higher affinity for the CAIX antigen, we anticipate that development of efficient SMCC products can make a valuable contribution in the field of cancer immunotherapy.

## Supporting information

Detailed experimental conditions

## 6. ASSOCIATED CONTENT

### Supporting Information

Detailed experimental conditions and additional data for the manuscript can be found here in the supplemental file.

## 7. Abbreviations

AAZ: acetazolamide
ADC: Antibody drug conjugate
CAIX: Carbonic anhydrase
IX IL2: Interleukin-2
LC-MS: Liquid Chromatography – Mass Spectrometry
RCC: Renal cell carcinoma
SMCC: Small molecule cytokine conjugate
SMDC: Small molecule drug conjugate
SPPS: solid phase peptide synthesis
SPR: Surface Plasmon Resonance
SrtA: Sortase A

## 8. ACKNOWLEDGEMENTS

Authors would like to thank Sarah Ducellier, Chiara Rondinini and Sheila Dakhel for the technical support during protein engineering and production. Financial support by the ETH Zürich, the Swiss National Science Foundation (grant number 310030_182003/1), the European Research Council (ERC) under the European Union’s Horizon 2020 research and innovation program (grant agreement 670603) is gratefully acknowledged.

## References

(1) Murer, P. and Neri, D. (2019) Antibody-cytokine fusion proteins: A novel class of biopharmaceuticals for the therapy of cancer and of chronic inflammation. N Biotechnol 52, 42–53.

(2) van den Heuvel, M. M., Verheij, M., Boshuizen, R., Belderbos, J., Dingemans, A.-M. C., De Ruysscher, D., Laurent, J., Tighe, R., Haanen, J. and Quaratino, S. (2015) NHS-IL2 combined with radiotherapy: preclinical rationale and phase Ib trial results in metastatic non-small cell lung cancer following first-line chemotherapy. J Transl Med 13, 32.

(3) Klein, C., Waldhauer, I., Nicolini, V. G., Freimoser-Grundschober, A., Nayak, T., Vugts, D. J., Dunn, C., Bolijn, M., Benz, J., Stihle, M., et al. (2017) Cergutuzumab amunaleukin (CEA-IL2v), a CEA-targeted IL-2 variant-based immunocytokine for combination cancer immunotherapy: Overcoming limitations of aldesleukin and conventional IL-2-based immunocytokines. Oncoimmunology 6, e1277306.

(4) Halin, C., Rondini, S., Nilsson, F., Berndt, A., Kosmehl, H., Zardi, L. and Neri, D. (2002) Enhancement of the antitumor activity of interleukin-12 by targeted delivery to neovasculature. Nat Biotech 20, 264–269.

(5) De Luca, R., Gouyou, B., Ongaro, T., Villa, A., Ziffels, B., Sannino, A., Buttinoni, G., Galeazzi, S., Mazzacuva, M. and Neri, D. (2019) A Novel Fully-Human Potency-Matched Dual Cytokine-Antibody Fusion Protein Targets Carbonic Anhydrase IX in Renal Cell Carcinomas. Front Oncol 9, 1228.

(6) Eigentler, T. K., Weide, B., de Braud, F., Spitaleri, G., Romanini, A., Pflugfelder, A., González-Iglesias, R., Tasciotti, A., Giovannoni, L., Schwager, K., et al. (2011) A dose-escalation and signal-generating study of the immunocytokine L19-IL2 in combination with dacarbazine for the therapy of patients with metastatic melanoma. Clin. Cancer Res. 17, 7732–7742.

(7) Catania, C., Maur, M., Berardi, R., Rocca, A., Giacomo, A. M. D., Spitaleri, G., Masini, C., Pierantoni, C., González-Iglesias, R., Zigon, G., et al. (2015) The tumor-targeting immunocytokine F16-IL2 in combination with doxorubicin: dose escalation in patients with advanced solid tumors and expansion into patients with metastatic breast cancer. Cell Adh Migr 9, 14–21.

(8) Osenga, K. L., Hank, J. A., Albertini, M. R., Gan, J., Sternberg, A. G., Eickhoff, J., Seeger, R. C., Matthay, K. K., Reynolds, C. P., Twist, C., et al. (2006) A phase I clinical trial of the hu14.18-IL2 (EMD 273063) as a treatment for children with refractory or recurrent neuroblastoma and melanoma: a study of the Children’s Oncology Group. Clin. Cancer Res. 12, 1750–1759.

(9) Marcucci, F., Bellone, M., Rumio, C. and Corti, A. (2013) Approaches to improve tumor accumulation and interactions between monoclonal antibodies and immune cells. MAbs 5, 34–46.

(10) List, T. and Neri, D. (2013) Immunocytokines: a review of molecules in clinical development for cancer therapy. Clin Pharmacol 5, 29–45.

(11) Hillier, S. M., Maresca, K. P., Lu, G., Merkin, R. D., Marquis, J. C., Zimmerman, C. N., Eckelman, W. C., Joyal, J. L. and Babich, J. W. (2013) 99mTc-Labeled Small-Molecule Inhibitors of Prostate-Specific Membrane Antigen for Molecular Imaging of Prostate Cancer. J Nucl Med, Society of Nuclear Medicine 54, 1369–1376.

(12) Vlahov, I. R. and Leamon, C. P. (2012) Engineering folate-drug conjugates to target cancer: from chemistry to clinic. Bioconjug. Chem. 23, 1357–1369.

(13) Ginj, M., Zhang, H., Waser, B., Cescato, R., Wild, D., Wang, X., Erchegyi, J., Rivier, J., Mäcke, H. R. and Reubi, J. C. (2006) Radiolabeled somatostatin receptor antagonists are preferable to agonists for in vivo peptide receptor targeting of tumors. PNAS, National Academy of Sciences 103, 16436–16441.

(14) Krall, N., Pretto, F., Mattarella, M., Müller, C. and Neri, D. (2016) A 99mTc-Labeled Ligand of Carbonic Anhydrase IX Selectively Targets Renal Cell Carcinoma In Vivo. J Nucl Med 57, 943–949.

(15) Cazzamalli, S., Corso, A. D. and Neri, D. (2017) Linker stability influences the anti-tumor activity of acetazolamide-drug conjugates for the therapy of renal cell carcinoma. J Control Release 246, 39–45.

(16) Lv, P.-C., Roy, J., Putt, K. S. and Low, P. S. (2017) Evaluation of Nonpeptidic Ligand Conjugates for the Treatment of Hypoxic and Carbonic Anhydrase IX-Expressing Cancers. Mol. Cancer Ther. 16, 453–460.

(17) Hilvo, M., Rafajová, M., Pastoreková, S., Pastorek, J. and Parkkila, S. (2004) Expression of carbonic anhydrase IX in mouse tissues. J. Histochem. Cytochem. 52, 1313–1322.

(18) Krall, N., Pretto, F. and Neri, D. (2014) A bivalent small molecule-drug conjugate directed against carbonic anhydrase IX can elicit complete tumour regression in mice. Chem. Sci. 5, 3640–3644.

(19) Cazzamalli, S., Dal Corso, A. and Neri, D. (2016) Acetazolamide Serves as Selective Delivery Vehicle for Dipeptide-Linked Drugs to Renal Cell Carcinoma. Mol. Cancer Ther. 15, 2926–2935.

(20) Cazzamalli, S., Ziffels, B., Widmayer, F., Murer, P., Pellegrini, G., Pretto, F., Wulhfard, S. and Neri, D. (2018) Enhanced Therapeutic Activity of Non-Internalizing Small-Molecule-Drug Conjugates Targeting Carbonic Anhydrase IX in Combination with Targeted Interleukin-2. Clin. Cancer Res. 24, 3656–3667.

(21) Kulterer, O. C., Pfaff, S., Wadsak, W., Garstka, N., Remzi, M., Vraka, C.soula, Nics, L., Bootz, F., Cazzamalli, S., Krall, N., et al. (2020) A microdosing study with 99mTc-PHC-102 for the SPECT / CT imaging of primary and metastatic lesions in renal cell carcinoma patients. J Nucl Med. doi: 10.2967/jnumed.120.245530.

(22) Rosenberg, S. A. (2014) IL-2: the first effective immunotherapy for human cancer. J. Immunol. 192, 5451–5458.

(23) Arenas-Ramirez, N., Woytschak, J. and Boyman, O. (2015) Interleukin-2: Biology, Design and Application. Trends Immunol. 36, 763–777.

(24) Chavez, A. R. de V., Buchser, W., Basse, P. H., Liang, X., Appleman, L. J., Maranchie, J. K., Zeh, H., de Vera, M. E. and Lotze, M. T. (2009) Pharmacologic administration of interleukin-2. Ann. N. Y. Acad. Sci. 1182, 14–27.

(25) Kalia, V., Sarkar, S., Subramaniam, S., Haining, W. N., Smith, K. A. and Ahmed, R. (2010) Prolonged interleukin-2Ralpha expression on virus-specific CD8+ T cells favors terminal-effector differentiation in vivo. Immunity 32, 91–103.

(26) Popp, M. W., Antos, J. M., Grotenbreg, G. M., Spooner, E. and Ploegh, H. L. (2007) Sortagging: a versatile method for protein labeling. Nat. Chem. Biol. 3, 707–708.

(27) Rashidian, M., Keliher, E. J., Dougan, M., Juras, P. K., Cavallari, M., Wojtkiewicz, G. R., Jacobsen, J. T., Edens, J. G., Tas, J. M. J., Victora, G., et al. (2015) Use of 18F-2-Fluorodeoxyglucose to Label Antibody Fragments for Immuno-Positron Emission Tomography of Pancreatic Cancer. ACS Cent Sci 1, 142–147.

(28) Witte, M. D., Wu, T., Guimaraes, C. P., Theile, C. S., Blom, A. E. M., Ingram, J. R., Li, Z., Kundrat, L., Goldberg, S. D. and Ploegh, H. L. (2015) Site-specific protein modification using immobilized sortase in batch and continuous-flow systems. Nat Protoc 10, 508–516.

(29) Bartels, L., Ploegh, H. L., Spits, H. and Wagner, K. (2019) Preparation of bispecific antibody-protein adducts by site-specific chemo-enzymatic conjugation. Methods 154, 93–101.

(30) Gébleux, R., Briendl, M., Grawunder, U. and Beerli, R. R. (2019) Sortase A Enzyme-Mediated Generation of Site-Specifically Conjugated Antibody-Drug Conjugates. Methods Mol. Biol. 2012, 1–13.

(31) Chen, L., Cohen, J., Song, X., Zhao, A., Ye, Z., Feulner, C. J., Doonan, P., Somers, W., Lin, L. and Chen, P. R. (2016) Improved variants of SrtA for site-specific conjugation on antibodies and proteins with high efficiency. Sci Rep 6, 31899.

(32) Frey, K., Schliemann, C., Schwager, K., Giavazzi, R., Johannsen, M. and Neri, D. (2010) The immunocytokine F8-IL2 improves the therapeutic performance of sunitinib in a mouse model of renal cell carcinoma. J. Urol. 184, 2540–2548.

(33) Wichert, M., Krall, N., Decurtins, W., Franzini, R. M., Pretto, F., Schneider, P., Neri, D. and Scheuermann, J. (2015) Dual-display of small molecules enables the discovery of ligand pairs and facilitates affinity maturation. Nature Chem 7, 241–249.

