## Supplementary material for "Sortase-mediated site-specific modification of interleukin-2 for the generation of a tumor targeting acetazolamide-cytokine conjugate": Detailed experimental conditions

1. Materials and Methods
   1.1 Small molecule generation

Gly-(Trt)-2-Chlorotrityl resin (600 mg) has been used for the synthesis of the peptide following the standard Fmoc-protocol. All amino acids were added sequentially according to the following sequence: Fmoc-Gly-OH , HOBt, HBTU, DIPEA, DMF, 1h, room temperature (r.t.) ; Fmoc-Thr-(tBu)-OH , HOBt, HBTU, DIPEA, DMF, 1h, r.t. ; Fmoc-Glu-(OtBu)-OH , HOBt, HBTU, DIPEA, DMF, 1h, r.t. ; Fmoc-Pro-OH , HOBt, HBTU, DIPEA, DMF, 1h, r.t. ; Fmoc-Leu-OH , HOBt, HBTU, DIPEA, DMF, 1h, r.t. ; 6-azido-hexanoic acid, HOBt, HBTU, DIPEA, DMF, 1h, r.t. ; N-(5-sulfamoyl-1,3,4-thiadiazol-2-yl)hex-5-ynamide, CuI, TBTA, DMF/THF 1:1, overnight, r.t. Cleavage of the peptide from the resin was performed using TFA (33 %), Triisopropylsilane (14%), mQ water (3%) and dichloromethane (50%). Compound (1) was purified using reverse-phase HPLC. Elution fractions were analyzed by mass spectrometry and compound (1) have been lyophilized overnight to obtain compound AAZ-LPETGG as a white powder (35 mg; 35.5 µmol; 7.2% yield). Quality of the pure material was analyzed by mass spectrometry.

1.2 Cloning of IL2 and sortase A

The genes encoding the human IL2 were generated using PCR. DNA of hIL2 with C125S mutation was used as template for the PCR. The linker (G_4_S)_3_ and the mutation T23S mutation were inserted by PCR with the use of primers containing the mutation (PCR1: Fw primer: GATCCTCCTGTTCCTCGTCGCTGTGGCTACAGGTGTGCACTCGGGTGGAGGCGGTTCAGGCGGAGGTGGCTCTGGCGGTGGCGGATCAGCACCTTCTTCA,Bw primer ACAGGCGGCCGCTTATCAATGGTGATGGTGGTGATGAGTCAGTGTTGAGA; PCR2: Fw primer TCCAGAAGCTTCCACCATGGGCTGGAGCCTGATCCTCCTGTTCCTCGTCG, Bw primer ACAGGCGGCCGCTTATCAATGGTGATGGTGGTGATGAGTCAGTGTTGAGA). The PCR product was then double digested with HindIII and NotI-HF restriction enzymes. The digested DNA fragments were ligated into pcDNA3.1 plasmid. Resulting DNA plasmids were amplified and used for mammalian cell transfection.

Whereas, Sortase A gene was amplify using a succession of two PCR in order to insert a Histidine tag for metal affinity purification. (PCR1: fw primer: GCCGCATATGATGCAGGCAAAACCGCAGATTCCGAA, bw primer: ATTAGCGGCCGCGTGATGGTGATGGTGATGTTCCAGT; PCR2: fw primer: ATTAGCATGCAAATTCTATTTCAAGGAGACAGTCATAATGCAGGCAAAACCGCAG, bw primer: GCCGGAATTCTTAGTGATGGTGATGGTGATGTTCCAGTTTAACTTCG). The PCR product was then double digested with SphI and EcoRI restriction enzymes. The digested DNA fragments were ligated into pUC119 plasmid. The resulting DNA plasmid was electroporated in TG1 bacteria for protein production.

1.3 Protein Expression, Purification and Characterization

Sortase A was produced by IPTG induction of electroporated TG1 when exponential phase of growth was reached. After 16h at 30°C 120 rpm bacteria were pelleted and resupanded in lysis buffer (50mM NaH2PO4, 300mM NaCl, 10mM Imidazole, 1mg/mL lysozyme pH 8) then sonicate for 5 minutes. Supernatant was collected by centrifugation (13000rpm for 15 minutes) and filtered (0.44um) prior Ninta affinity chromatography (GE healthcare) following the protocol provided by the supplier. G4S3-IL2 and the anti-IL2-Fab were expressed using transient gene expression in CHO cells. (Hacker et al., 2013) For 1 ml of production, 4 × 10^6^ cells were collected by centrifugation and resuspended in 1 mL ProCHO4 (LONZA) supplemented with 4 mM Ultraglutamine (Lonza). Per million cells, 0.625 ug of plasmid DNAs followed by 2.5 ug polyethylene imine (PEI; 1 mg/mL solution in water at pH 7.0, Polysciences) were added to the cells and gently mixed. The transfected cultures were incubated in a shaker incubator at 31°C in an incubator with 5% CO2 atmosphere shaking at 120rpm for 6 days. The His-tagged proteins were purified by affinity chromatography using Ninta affinity chromatography (GE Healthcare) following the protocol provided by the supplier. Proteins were dialyzed overnight at 4°C against PBS prior utilization. For the generation of the affinity chromatography against IL2, the anti-IL2 antibody produced inhouse was coupled to CNBr Activated Sepharose® 4 Fast Flow (GE Healthcare) following the manufacturer instructions. After the enzymatic coupling reaction AAZ-IL2 was purified in homogeneity using the anti-IL2 chromatography following Protein A purification described elsewhere (Ongaro T et al., J Biotechnol. 2019.) Elution were pooled and dialyzed against PBS. Purified proteins were characterized for their size, homogeneity and purity by SDS-PAGE, Size exclusion chromatography and LC-MS, respectively. For SDS-PAGE analysis proteins were run under reducing and non-reducing conditions on 10 or 12% acrylamide gels (Invitrogen) and stained using Coomassie blue. Size-exclusion chromatography was performed on an ÄKTA FPLC system using a Superdex 75 increase 10/300GL column (Amersham Biosciences). For the LC-MS, proteins were injected on a Waters Xevo G2XS Qtof instrument (ESI-ToF-MS) coupled to a Waters Acquity UPLC H-Class System using a 2.1 × 50 mm Acquity BEH300 C4 1.7 µm column (Waters). 0.1% FA in water (solvent A) and 0.1% FA in MeCN (solvent B) were used as mobile phase at a flow rate of 0.4 mL/min. Gradient was programmed as follows: after 1.5 min isocratic with 95% solvent A, stepwise change from 95% solvent A to 95% solvent B in 4.5 min (10% increase every 0.5 min), back to 95% solvent A in 0.5 min, linearly to 95% solvent B and back to 95% solvent A in 2.25 min (last step repeated twice). AAZ-IL2 binding affinity was evaluated by Surface plasmon resonance using CM5 chip (GE Healthcare) coated with recombinant hCAIX. Sample at a concentration of 500nM was injected on a BIAcore X100 instrument (GE Healthcare). For fluorescent labeling of AAZ-IL2 and G4S3-IL2, proteins were dialyzed against 0.1M Sodium Carbonate pH 9 over night at 4°C. 25uL of 1mg/mL of FITC (Acros organics) was added to 1 mg of protein for 16h at 4°C under gentle agitation. FITC labeled protein were purified over size exclusion chromatograpy (PD10, GE Healthcare).

1.4 Key Experimental procedure

For the conjugation of the small molecule cytokine conjugate, 100 µM of the purified “sorttaged” small molecule (AAZ-LPETGG) was incubated at 4°C for 3h with 5 µM of Sortase A and 5µM of “sorttaged” IL2 cytokine ((G4S)3-IL2) under gentle agitation. Once the reaction was complete, the final product was purified to homogeneity via affinity chromatography. Briefly, the crude was incubated 2 hours at room temperature with an anti-IL2 resin. The resin was then washed with Buffer A (100mM NaCl,0.5mM EDTA, 0.1%tween, PBS) and Buffer B (500mM NaCl,0.5mM EDTA, PBS). Elution was performed with 0.1M glycine pH 3 and fractions containing the AAZ-IL2 product were pooled and dialyzed overnight against PBS (100mM NaCl 30mM Na_2_HPO_4_, 20mM NaH_2_PO4). Pure AAZ-IL2 was used for SPR analysis, iodinated biodistribution and FITC labeling for immunofluorescence biodistribution. Detailed experimental procedure is presented as part of the supporting information.

1.1 Tumor targeting via radiolabeling and immunofluorescence

In vivo experiments on tumor models were performed under a project license granted by the cantonal veterinary office (ZH004/18) in agreement with Swiss regulations. For SKRC52 were grown in RPMI (Life Technologies) supplemented with 10% FBS at 37 °C and 5% CO2, cells were detached using Trypsin-EDTA 0.05% (Invitrogen). Female athymic Balb/c nu/nu mice (8 weeks old, Janvier) were injected on the right flank with 5×10^6^ SKRC52 cells. As described previously (Pfaffen et al., 2010), *In vivo* targeting was evaluated by biodistribution analysis. Proteins were radioiodinated with ^125^I and Chloramine T hydrate and purified on a PD10 column (GE Healthcare). ~10 ug of Radiolabeled proteins were injected intravenously. Mice were sacrificed at different timepoints (1h, 3h, 6h, 24h). Organs were weighed and radioactivity was counted using a Packard Cobra gamma counter (Packard, Meriden, CT, USA). Reported values are in injected dose per gram of tissue (%ID/g). For immunofluorescence targeting, mice were injected intravenously with Fluorescent derivatives. After 2h, mice were sacrificed and organs were embedded and frozen into OCT. Frozen sections (8um) were fixed by 4% of Formalin and blocked with 20% fetal bovine serum in PBS. Detection of the test article was performed using rabbit anti-FITC igG (Biorad) followed by anti-rabbit IgG Alexa 488 (biotum) and counter stained with DAPI.
