## Supplementary material for "Sortase-mediated site-specific modification of interleukin-2 for the generation of a tumor targeting acetazolamide-cytokine conjugate": Figure S1

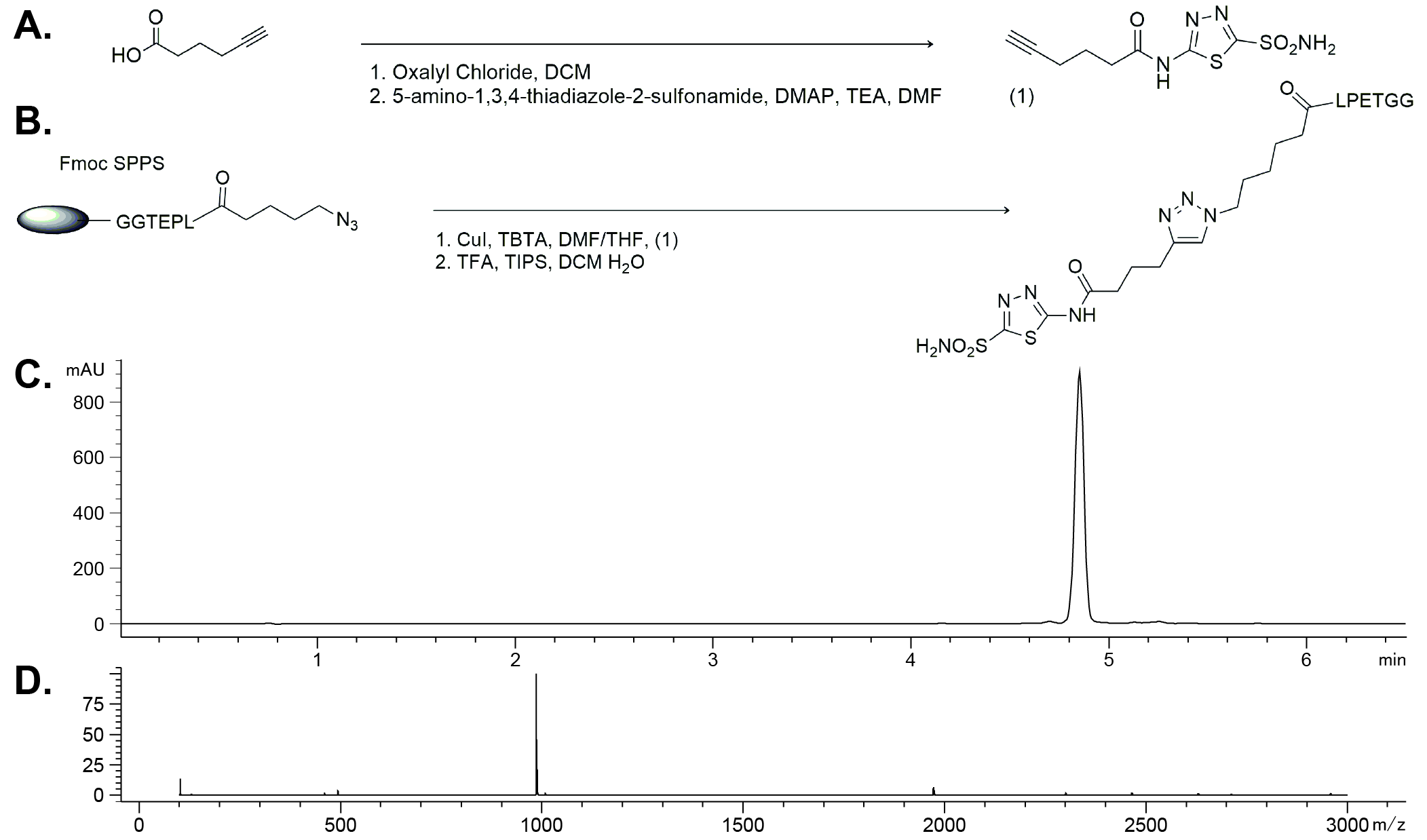

**Figure S1: AAZ-LPETGG purity characterization after chemical purification.** (A) Schematic representation of compound 1. (B) Schematic representation of SPPS production of AAZ-LPETGG. (C) OD260 of the LC-MS of AAZ-LPETGG. (D) m/z of AAZ-LPETGG after LC-MS quality control.
