## Supplementary material for "Sortase-mediated site-specific modification of interleukin-2 for the generation of a tumor targeting acetazolamide-cytokine conjugate": Figure S2

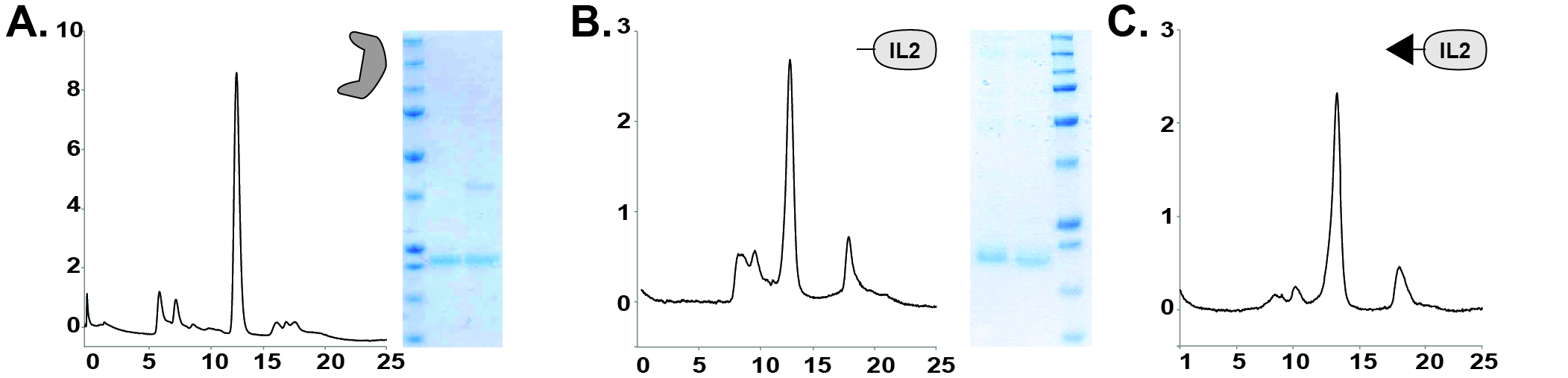

**Figure S2: Proteins produced exhibit a good quality after production and purification.** Size exclusion chromatography and SDS Page of (A) Sortase A (B) G4S3-IL2 after affinity chromatography purification. Size exclusion chromatography of (C) AAZ-LPETGG after affinity chromatography purification.
